# Spatial organization of myofibroblastic and complement-secreting CAFs in neuroendocrine tumors

**DOI:** 10.64898/2026.02.26.708188

**Authors:** Helvijs Niedra, Marta L. Springe, Katrine Tiltina, Raitis Peculis, Rihards Saksis, Jurijs Nazarovs, Arturs Ozolins, Sofija Vilisova, Natalja Senterjakova, Aija Gerina, Ilze Konrade, Aldis Pukitis, Vita Rovite

**Affiliations:** Department of Molecular and Functional Genomics, Latvian Biomedical Research and Study Centre, Riga, LV-1067, Latvia; Pauls Stradins Clinical University Hospital, Riga, LV-1002, Latvia; Department of Pathology, Riga Stradins University, Riga, LV-1007, Latvia; Department of Endocrinology, Riga East Clinical University Hospital, Riga, LV-1079, Latvia; Department of internal diseases, Riga Stradins University, Riga, LV-1007, Latvia

## Abstract

Neuroendocrine tumors are graded and classified largely by tumor cell-intrinsic features, yet the stromal microenvironment remains poorly defined across anatomical sites. We applied near single-cell spatial transcriptomics augmented with cell segmentation to eight treatment-naive neuroendocrine tumor primary tissues from pancreas, colon, appendix, and bile duct to build a spatially resolved stromal reference. Integration of stromal-enriched cell polygons identified ten transcriptional states shared across tumors, with a minority of niche-restricted clusters mapping to tumor/stroma and stroma/non-tumor boundaries. Within the shared fibroblast compartment, program scoring resolved four cancer-associated fibroblast states. Myofibroblastic and complement-secretory states dominated across samples, whereas inflammatory and antigen-presenting programs were consistently detected but weaker. Spatial mapping in desmoplastic tumors showed myofibroblastic fibroblasts enriched in collagen-dense regions, while complement-secretory fibroblasts localized preferentially to tumor-adjacent stromal interfaces. Pseudobulk differential expression and gene set enrichment analyses supported extracellular matrix remodeling in myofibroblastic fibroblasts and complement cascade activation in complement-secretory fibroblasts. Together, these findings demonstrate that anatomically distinct neuroendocrine tumors share a conserved yet spatially segregated stromal architecture, characterized by dominant matrix-producing and complement-enriched fibroblast states.

## 1. Introduction

Neuroendocrine (NE) neoplasms are a rare and heterogeneous subgroup of epithelial neoplasms arising from diffuse neuroendocrine cells (Rindi *et al*, 2022). Although they can occur throughout the body, most originate in the digestive tract and lungs. In contrast to poorly differentiated neuroendocrine carcinomas, well-differentiated neuroendocrine tumors (NETs) typically retain organ-specific lineage features, show variable hormonal activity, and often follow a more indolent clinical course (Rindi *et al*, 2022). Clinically, NETs are stratified into three grades based on proliferative activity (mitotic count and Ki-67 index), with increasing grade reflecting progressively more aggressive behavior (Rindi *et al*, 2022). While grading is prognostically informative, it does not fully capture the biological diversity that underlies differences in clinical course within and across anatomical sites. In routine practice, chromogranin A (CHGA) and synaptophysin (SYP) remain widely used markers to support diagnosis and disease monitoring (Tomita, 2020; Rindi *et al*, 2022), yet these lineage markers provide limited insight into whether NETs share conserved transcriptional, chromatin and stromal cell transcriptional programs across sites, or instead follow predominantly anatomy-defined molecular trajectories.

Among gastroenteropancreatic NETs (GEP-NETs), pancreatic neuroendocrine tumors (PanNETs) represent a prominent subgroup (Rindi *et al*, 2022). PanNETs frequently harbor alterations in chromatin remodeling genes, most notably ATRX and DAXX, which are linked to alternative lengthening of telomeres and distinct clinical outcomes (Yasunaga *et al*, 2024; van T Veld *et al*, 2025). However, these alterations are not universal and are insufficient for comprehensive tumor stratification, particularly in settings where improved molecular resolution could inform risk assessment earlier in disease. More recently, lineage-associated transcription factors including ISL1 and CDX2, as well as ARX and PDX1, marking α-cell and β-cell lineage commitment, have been highlighted as predictors of tumor phenotype, functional status, and invasive potential (Lakis *et al*, 2021; Moser *et al*, 2024). Collectively, these approaches refine tumor classification but remain primarily tumor cell centric.

Beyond tumor-intrinsic programs, the tumor microenvironment (TME) is increasingly recognized as a major determinant of NET progression and therapeutic response (Cives *et al*, 2019; Chen *et al*, 2024). The stromal compartment - comprising fibroblasts, vasculature, extracellular matrix, myeloid cells, and other immune components - can either support or constrain tumor growth depending on its activation state (Xu *et al*, 2022). Consistent with this, recent studies report clinically relevant immune-stromal features in NETs, including tumor-associated macrophages and elevated PD-1 expression in pancreatic neuroendocrine neoplasms (Xu *et al*, 2022), as well as prominent α-smooth muscle actin (α-SMA)-positive stromal cell populations indicative of activated stroma PanNETs (Niedra *et al*, 2025)

A central stromal component of most solid tumors is the cancer-associated fibroblast (CAF), which is functionally heterogeneous and includes antigen-presenting CAFs (apCAFs), inflammatory CAFs (iCAFs), myofibroblastic CAFs (myCAFs), as well as emerging complement-secretory (csCAFs) and intermediate phenotypes (Chen *et al*, 2026; Yang *et al*, 2023). These states differentially regulate extracellular matrix remodeling, cytokine secretion, immune modulation, and therapeutic resistance - where myCAFs are frequently linked to matrix densification and impaired drug penetration, while iCAFs are more closely associated with cytokine-driven inflammatory signaling (Yang *et al*, 2023). Across tumor types, CAF programs have been repeatedly associated with disease progression and therapy resistance (Hu *et al*, 2019; Linares *et al*, 2020; Yasuda *et al*, 2021), and multiple CAF states can coexist within the same tumor forming distinguishable tumor-stroma interactomes, contributing to substantial intratumoral heterogeneity (Forsthuber *et al*, 2024; Chen *et al*, 2026). Importantly, CAF function is strongly shaped by spatial context, particularly their proximity to tumor cells, vasculature, and immune infiltrates (Chen *et al*, 2026). Despite these advances, the distribution, transcriptional features, and spatial interactions of CAF subpopulations in NETs remain incompletely defined.

Recent advances in spatial transcriptomics enable high-resolution mapping of gene expression in intact tissue, allowing simultaneous interrogation of tumor cells and stromal and immune compartments within preserved architecture (Oliveira *et al*, 2025). This provides a framework to link tumor-intrinsic programs with spatially organized microenvironmental states and to resolve how stromal subtypes are arranged within tumor niches. Accordingly, in this study, using Visium HD spatial transcriprtomics method (Oliveira *et al*, 2025), we profiled eight well-differentiated NETs: five PanNETs, one appendiceal (aNET), one colorectal (cNET), and one biliary (bNET) neuroendocrine tumor. We aimed to (i) characterize stromal heterogeneity across anatomically distinct NETs and (ii) map the spatial organization of four CAF programs (myCAFs, csCAFs, iCAFs, apCAFs). By integrating spatial profiles across five NET entities, we seek to establish a stromal reference for GEP-NETs and to define conserved versus context-specific microenvironmental features with potential relevance to NET classification and future biomarker and therapeutic strategies.

## 2. Materials and methods

### 2.1. Study cohort

The study cohort comprised eight retrospective, treatment-naive patients (two male, six female) diagnosed with GEP-NETs, including five PanNETs, one aNET, one cNET, and one bNET (Table 1). Patient age at diagnosis ranged from 23 to 67 years (median - 56 years). At the time of diagnosis, none of the patients showed distant metastatic disease; however, two patients (NET078 and NET017) presented with regional lymph node involvement. Accordingly, only the formalin-fixed, paraffin-embedded (FFPE) primary tumor tissue was obtained for this study. All study procedures were conducted in accordance with the principles of the Declaration of Helsinki and were approved by the Central Medical Ethics Committee of Latvia (approval protocol No. 01-29.1.2/1908). Patients were recruited from Pauls Stradins Clinical University Hospital and Riga East Clinical University Hospital. Written informed consent was obtained from all participants prior to inclusion in the study.

**Table 1.** Clinical characteristics of patients recruited in the study.

| Patient ID | Year | Sex | Age of diagnosis | NET primary localization | TNM | Grade |
| --- | --- | --- | --- | --- | --- | --- |
| NET040 | 2022 | F | 57 | PanNET | pT2N0M0R0 | G1 |
| NET078 | 2022 | M | 57 | PanNET | pT2N1M0Rx | G1 |
| NET084 | 2022 | F | 41 | PanNET | pT2N0M0Rx | G1 |
| NET109 | 2024 | F | 54 | PanNET | pT2N0M0R0 | G3 |
| NET111 | 2024 | M | 67 | PanNET | pT1N0M0Rx | G1 |
| NET017 | 2019 | F | 63 | cNET | pT3N1M0R0 | G1 |
| NET047 | 2022 | F | 23 | aNET | pT1N1M0R0 | G1 |
| NET022 | 2021 | F | 42 | bNET | pT1N0M0R0 | G3 |

Abbreviations: G1 - grade 1; G3 - grade 3; PanNET - pancreatic neuroendocrine tumor; cNET - colorectal neuroendocrine tumor; aNET - appendiceal neuroendocrine tumor; bNET - biliary neuroendocrine tumor.

### 2.2 FFPE sample quality control for spatial profiling

Upon collection, FFPE tissue blocks were sectioned to obtain two 10 µm-thick slices, which were immediately stored at −80 °C until RNA extraction. Total RNA was extracted from FFPE tissue sections using the RNeasy FFPE Kit (Qiagen). RNA concentration was measured using a Qubit 4 Fluorometer with the Qubit RNA High Sensitivity assay (Thermo Fisher Scientific). RNA quality was assessed using the TapeStation 4200 with High Sensitivity RNA ScreenTape (Agilent). All samples met the minimum quality requirements for Visium HD analysis, with DV200 values exceeding 30% and ranging from 47% to 80%.

### 2.3 Sample preparation for spatial transcriptomics analysis

Visium HD spatial transcriptomics library preparation and sequencing were performed as a commercial service by the Innovative Medical Center at the Institute of Human Genetics, Polish Academy of Sciences (Poznań, Poland), following the manufacturer’s recommended protocols for FFPE tissue. FFPE blocks were stored at 4 °C prior to processing. Tissue sections (5 µm thickness) were prepared using a microtome and initially mounted on standard glass slides for histological quality control, imaging, and selection of representative regions. Hematoxylin and eosin (H&E) staining and imaging were performed in accordance with the Visium HD FFPE Tissue Preparation guidelines. Spatial gene expression profiling was carried out using the 10x Genomics Visium HD platform and whole-transcriptome probe panels targeting the human transcriptome. Probe hybridization and ligation were performed directly within the tissue sections. Slides were subsequently processed using the Visium CytAssist instrument to enable probe release and capture onto spatially barcoded oligonucleotides on Visium HD slides. Following probe capture, slides were processed for Illumina-based next-generation sequencing library preparation. Libraries were generated separately for each tissue sample, and library concentration and quality were assessed prior to pooling. Pooled libraries were sequenced on an Illumina NextSeq 2000 platform using a 1.8 B flow cell with a paired-end read configuration of 43-10-10-50 (Read 1-i7-i5-Read 2) and a 1% PhiX spike-in, in accordance with the Visium HD protocol.

### 2.4 Data preprocessing and cell segmentation

Raw sequencing data (FASTQ files) were processed using the Space Ranger (v4.0) pipeline to align sequencing reads and map spatial gene expression to high-resolution hematoxylin and eosin (H&E) tissue images. To achieve near single-cell spatial resolution, custom cell segmentation masks were generated independently for each sample. Nuclei segmentation followed by cytoplasmic expansion was performed using QuPath (v0.6.0) (Bankhead *et al*, 2017) in combination with the StarDist extension (v0.6.0) (Schmidt *et al*, 2018), applying the developer-provided model (*he_heavy_augment*) optimized for H&E-stained tissue. Default segmentation parameters were manually adjusted to account for tissue-specific differences in morphology and staining intensity. The resulting segmentation masks were supplied to Space Ranger to enable aggregation of 2 µm spatial bins into individually segmented cells / spatial polygons, generating a cell boundary-aware spatial transcriptomic dataset.

### 2.5 Initial identification of stromal cell compartments

Downstream analyses were performed in R using Seurat (v5) (Butler *et al*, 2018), installed from the satijalab GitHub repositories (ref: spaceranger-4.0) to ensure compatibility with Space Ranger v4.0 segmentation outputs. At this stage, no additional quality control or filtering steps were applied. Initial clustering results generated by the Space Ranger v4.0 pipeline were imported into Seurat objects and used to guide identification of stromal compartments within each tumor. Gene expression values were normalized using the standard Seurat LogNormalize method with a scale factor corresponding to the median total transcript count per segmented cell polygon. To characterize the transcriptional identity of Space Ranger-derived clusters, differential gene expression analysis was performed using the Wilcoxon rank-sum test. For each cluster, the top 100 upregulated genes (ranked by percentage difference) were selected and subjected to cell-type enrichment analysis using EnrichR web server (Kuleshov *et al*, 2016). Enrichment results with the PanglaoDB (Franzén *et al*, 2019) cell type gene sets database was integrated with spatial expression patterns in 10x genomics Loupe browser v9.0 to guide identification of stromal-enriched clusters for downstream analyses.

### 2.6 Program score-based quality control and filtering of stroma-enriched subsets

During initial sample inspection using Loupe Browser (v9.0), all samples were observed to exhibit varying degrees of lateral RNA diffusion, manifested by detectable expression of neuroendocrine markers (e.g., INS, CHGA, CHGB) and secretory epithelial markers (e.g., CPA1, CPB1) within stromal-assigned segments. To ensure that downstream stromal clustering analyses were not driven by highly expressed diffuse ambient transcripts due to lateral RNA-diffusion (Liang *et al*, 2023) or residual epithelial cells insufficiently separated by initial Space Ranger clustering. We devised an additional gene program score-based quality control method to remove cell polygons that were highly indicative of mixed cells or residual epithelial cells.

Accordingly, curated gene programs were constructed to define expected major TME-associated stromal, and immune cell lineages, including fibroblasts, endothelial cells, pericytes, smooth muscle cells, myeloid cells, B/plasma cells, T/NK cells, and neural/Schwann cells (Supplementary Table 1). Gene selection was guided by publicly available annotations from the Human Protein Atlas (Thul & Lindskog, 2018) and manual visualization of expression in Loupe browser; only the expressed candidate genes were selected to avoid oversized gene lists. In parallel, negative control programs were defined to capture tumor-derived neuroendocrine signatures, epithelial lineages (ductal, exocrine, and intestinal), and red blood cell-associated transcripts. Expression scores for each program were calculated using Seurat v5 inbuilt AddModuleScore feature. However, to better account for sample-specific differences in expression scale, program scores were transformed into percentile ranks. Percentile-based thresholds were subsequently applied to flag and exclude polygons exhibiting either mixed stromal/immune lineage programs or strong enrichment of epithelial, neuroendocrine, or red blood cell-associated transcriptional signatures.

Following the removal of polygons that exhibited strong mixed lineage or epithelial signal, standard quality control (QC) metrics were also applied. To mitigate the impact of potential segmentation artifacts resulting in multi-cellular doublets, cells exceeding the 99th percentile of either total transcript counts (nCounts) or detected features (nFeatures) were excluded. Furthermore, the percentage of mitochondrial gene expression was calculated for each sample; cells exhibiting extreme mitochondrial signals (above the 99th percentile) were removed to prevent the inclusion of dying or stressed cells in downstream analyses.

### 2.7 Stromal subset integration and clustering in integrated space

Stromal-enriched subsets from all eight samples were integrated and jointly analyzed in R using Seurat v5 to enable cross-sample comparison of stromal cell states. For each sample, normalized expression was computed using log-normalization with a common scale factor defined as the median of per-sample median library sizes (median UMI counts per sample stromal subset). Highly variable features were identified per sample using the “vst” method (2,000 genes), and a combined set of 3’000 integration features was selected across samples. For each sample, data was scaled and principal component analysis (PCA) was performed using the combined integration features and up to 30 PCs. Anchor-based integration was conducted using reciprocal PCA (RPCA) with 15 dimensions to identify shared stromal states (anchors) across samples. Integrated expression values were generated using the first 10 integration dimensions (decision based on per sample elbow plots). The integrated assay was subsequently re-scaled, and PCA was rerun prior to graph-based clustering. A shared nearest-neighbor (SNN) graph was constructed using the first 10 PCs again and with k.param = 100, and clusters were identified using the Leiden algorithm (resolution = 0.4). Two-dimensional embeddings were generated using UMAP based on the first 10 PCs and n.neighbors = 100. Lastly, cluster marker genes were identified in the non-integrated assay using Seurat’s Wilcoxon rank-sum test. Followingly, the top 100 marker genes from each cluster were once again used to derive identity of each cluster via EnrichR web server.

### 2.8 Cancer associated fibroblast subtype identification in shared fibroblast-like cell cluster

To characterize cancer-associated fibroblast (CAF) subtypes/states, the fibroblast-like stromal cluster (Cluster 1) identified in the integrated analysis (Section 1.7) was subsetted for downstream CAF subtyping. CAF state programs representing myCAFs, iCAFs, apCAFs, and csCAFs were defined based on published CAF marker sets (Zhu *et al*, 2025; Elhashani *et al*, 2024; Mori *et al*, 2024; Cui Zhou *et al*, 2022; Chen *et al*, 2021a; Zhang *et al*, 2025; Tao *et al*, 2025; Melchionna *et al*, 2023; Chen *et al*, 2021b, 2026; Yang *et al*, 2023). As the Visium HD Human Probe Set v2 did not include classical MHC-II genes, the gene set for apCAFs was supplemented using STRING protein-protein interaction network resources (v12.0) to identify genes involved in MHC-II complex transcription and antigen-processing pathways (Szklarczyk *et al*, 2023). Final gene lists used for CAF subtyping are provided in Supplementary Table 1.

Program scores for each CAF state were computed per sample using Seurat’s AddModuleScore in the non-integrated assay. CAF state assignment was performed using a competitive scoring framework designed to avoid ambiguous calls. Briefly, CAF program scores were standardized within each sample (z-score transformation) to account for sample-specific shifts in scoring scale. For each segmented unit, the highest-scoring CAF program was selected as the provisional state (“winner”) and required to exceed the second-highest program by a minimum margin (Δz ≥ 0.25) and to meet a minimum standardized strength threshold (z ≥ 0.25). To reduce spurious classifications, two additional safeguards were applied: 1) the raw module score of the winning program was required to be ≥ 0.05 to prevent inflated z-scores if a specific subtype contained multitude of near-zero values; and 2) gene-level evidence was required, defined as detection (raw counts > 0) of at least two genes from the corresponding program.

Lastly, to validate CAF subtype assignments and to derive cohort-shared CAF subtype marker genes a pseudobulk-style differential expression analysis, in which counts were aggregated per subtype per sample and analyzed using DESeq2 (Love *et al*, 2014), was performed and followed by gene set enrichment analysis using FGSEA package (Korotkevich & Sukhov, 2021) with Reactome (Milacic *et al*, 2024) and MSigDB Hallmark gene sets (Liberzon *et al*, 2015).

## 3. Results

### 3.1 Cohort-wide spatial transcriptomic profiling GEP-NET tissues

Spatially resolved transcriptomic profiling was performed on eight GEP-NET primary tumor samples (Figure 1A). Across all samples, the median unique molecular identifier (UMI) count per segmented cell polygon was 141 (range: 62 - 494), and the median number of detected genes per polygon was 112. The median polygon area ranged from 60 to 176 µm² (overall median: 134 µm²). Detailed sequencing and segmentation metrics, including RNA quality indicators (DV200), are provided in Supplementary Table 2.

**Figure 1.**
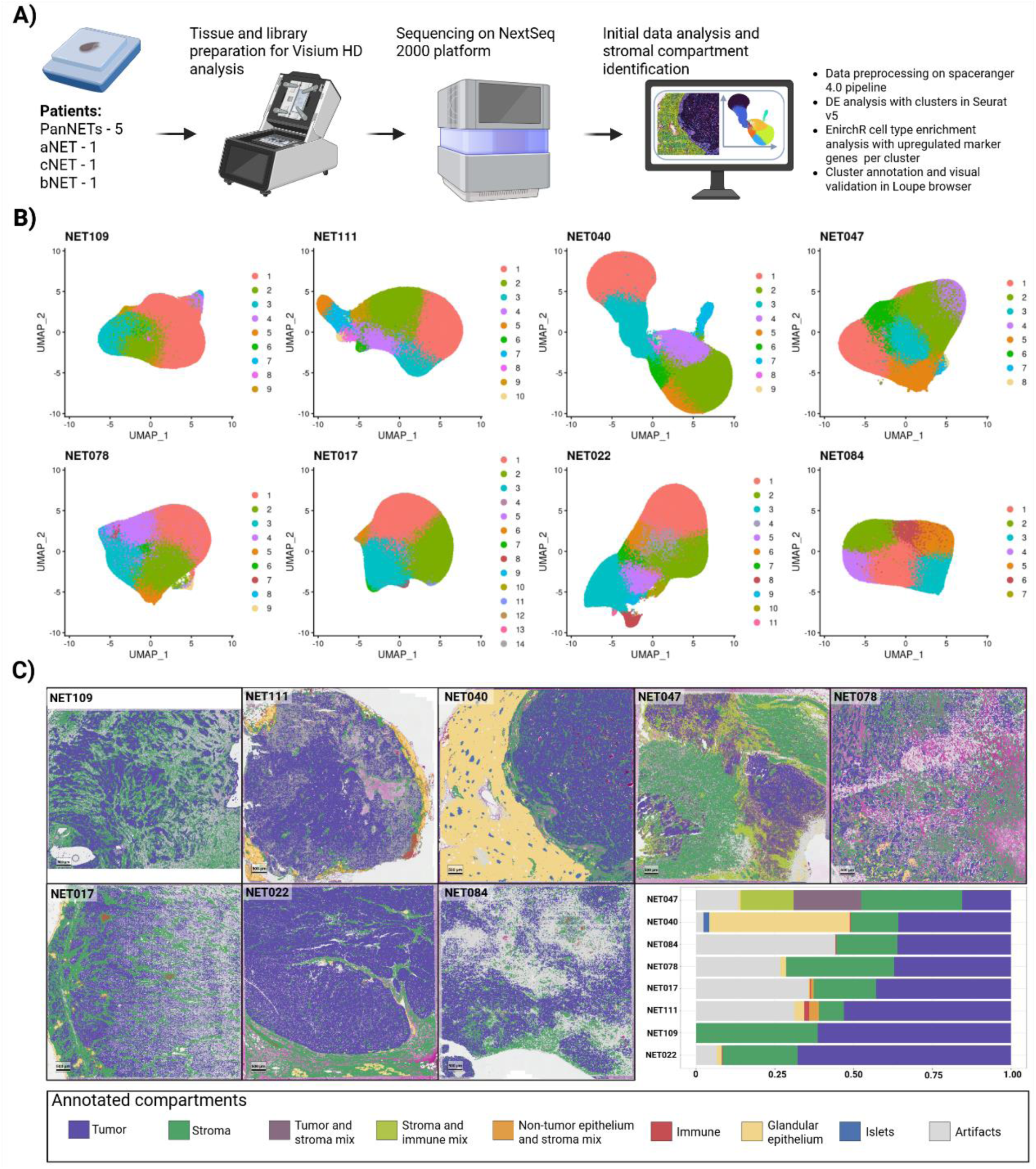
Spatial transcriptomic profiling of gastroenteropancreatic neuroendocrine tumors using Visium HD. **(A)** Overview of the experimental workflow. Eight tumor samples (PanNET, n = 5; aNET, n = 1; cNET, n = 1; bNET, n = 1) were processed using the 10x Genomics Visium HD platform. Libraries were sequenced on a NextSeq 2000 instrument. Raw data were processed using the Space Ranger 4.0 pipeline, followed by downstream visualization of Space Ranger pipeline generated clusters and differential expression analysis in Seurat v5. This was followed by cell type enrichment analysis using EnrichR. Resulting preliminary cluster annotations were validated by visual inspection in Loupe Browser. **(B)** UMAP representations of segmented cell polygons for each tumor sample. Cells are colored according to initial Space Ranger cluster assignments. UMAP coordinates were standardized across samples to enable direct visual comparison of cluster structure and relative transcriptional heterogeneity. **(C)** Spatial mapping of annotated cellular compartments overlaid on H&E images for each tumor section. Major compartments include tumor, stroma, immune, glandular epithelium, islets, mixed regions, and artifacts (clusters with no representative markers). The bar plot summarizes proportional compartment composition across samples. For sample NET109, minor peripheral tissue loss occurred during image stitching prior to CytAssist alignment, resulting in reduced coverage of the H&E image relative to the capture area. Spatial segmentation was therefore restricted to the imaged region.

Initial unsupervised clustering using the Space Ranger v4.0 pipeline identified major tissue compartments within each sample (Figure 1B). Cluster annotation, based on differential expression analysis and EnrichR cell-type enrichment, revealed tumor cells, stromal cells, immune infiltrates, and adjacent normal structures such as endocrine islets or glandular epithelium. The relative abundance of these compartments varied substantially between samples (Figure 1C), reflecting both biological heterogeneity and differences in tissue composition within the selected capture areas. The proportion of stromal segments within profiled regions ranged from 8% to 38% (median 22%). The highest stromal proportions were observed in NET109 (38%), NET078 (34%), and NET047 (32%). Adjacent non-tumorous tissue was predominantly detected in NET040, where morphologically normal exocrine pancreas and islet compartments constituted approximately 46% of the capture area. Corresponding high-resolution H&E images of the spatially profiled areas are available in Supplementary Figure 1. In the remaining samples, only a minor fraction (<3%) of profiled regions contained non-neoplastic epithelial cells. Differentially expressed genes used for cluster annotation in each sample are provided in Supplementary Table 3.

### 3.2 Per sample characterization of stromal compartment architecture

Marked heterogeneity in stromal architecture was observed across tumors. The most desmoplastic samples were the pancreatic NETs NET109 and NET078, in which unsupervised clustering resolved multiple transcriptionally distinct stromal subpopulations. In NET109, Clusters 2 and 3 were enriched for fibroblast and pancreatic stellate cell markers, Clusters 6 and 9 corresponded to vascular and smooth muscle compartments, Cluster 5 displayed neural/Schwann signatures, and Cluster 8 exhibited a mixed immune-fibroblast-neural profile. Similarly, NET078 contained two transcriptionally distinct fibroblast populations (Clusters 3 and 4), while additional clusters corresponded to vascular and mural cell compartments (Figure 2A-C). Spatial mapping revealed distinct localization patterns of fibroblast populations in both NET109 and NET078: one population was situated immediately adjacent to tumor nests, whereas the second localized deeper within extracellular matrix-rich stromal regions. Differential expression analysis specifically comparing tumor-adjacent and tumor-distant fibroblast clusters demonstrated that the tumor-proximal fibroblast population exhibited increased expression of neuroendocrine-related genes, most notably *CHGA* and *CHGB* (Supplementary Table 4). Histologically, NET078 also exhibited extensive stromal hyalinization, characterized by intense eosinophilic staining on H&E (Supplementary Figure 1). This feature likely reduced the density of detectable stromal nuclei and may have contributed to underestimation of stromal abundance in segmentation-based quantification.

**Figure 2.**
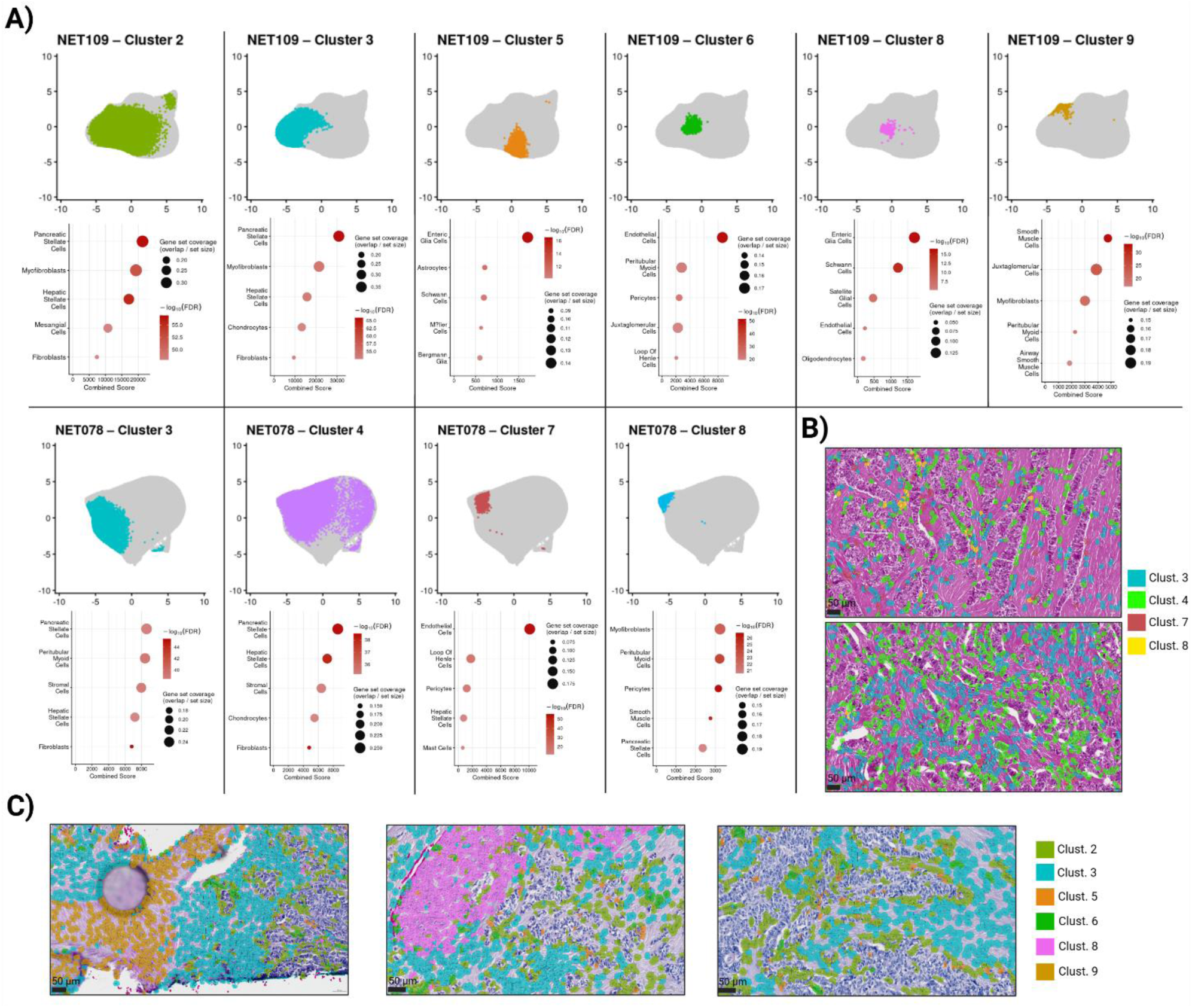
Pre-integration characterization of stromal compartment heterogeneity. **(A)** UMAP representations of segmented cell polygons transcriptomic profiles from two representative pancreatic neuroendocrine tumors (NET109 and NET078) showing stromal subclusters identified by Space Ranger prior to dataset integration. Clusters with stromal transcriptional signatures are highlighted individually, and their corresponding cell type enrichment profiles based on top 100 upregulated DEGs subjected to enrichR overrepresentation analysis with PanglaoDB Augmented 2021 database are shown below. Dot size indicates gene set coverage (overlap / gene set size), and color intensity represents -log10(FDR). These results demonstrate transcriptionally distinct stromal subpopulations within individual tumors prior to integration. (B) Spatial mapping of selected stromal clusters from NET078 overlaid on H&E images, illustrating distinct spatial localization patterns within the tumor microenvironment. (C) Spatial mapping of stromal clusters in NET109, further confirming the presence of multiple transcriptionally and spatially coherent stromal compartments.

NET047 (appendiceal NET) displayed the most complex stromal architecture. Histologically, tumor cells invaded adjacent smooth muscle tissue, forming a fibromuscular stromal network interspersed with tumor nests (Supplementary Figure 1). Consistent with this morphology, Cluster 1 exhibited SMC-like transcriptional signatures representing invaded smooth muscle tissue. However, due to extensive spatial intermixing of tumor, stromal, and immune compartments, unsupervised clustering did not fully resolve discrete populations. Cluster 2 (21% of cell polygons) displayed mixed tumor-stromal transcriptional profiles, while Cluster 3 (17%) exhibited stromal-immune signatures (Figure 1B; Supplementary Table 3). In the remaining samples (NET022, NET084, NET017, NET111, NET040), the stroma exhibited a more classical fibrovascular architecture. In NET040 and NET111, only a single dominant stromal cluster was identified (Cluster 3 and Cluster 4, respectively). These clusters expressed composite fibrovascular signatures, including extracellular matrix genes (COL3A1, COL1A1, COL1A2, DCN, BGN, LUM), endothelial markers (VWF, PECAM1), mural genes (RGS5, NOTCH3, MYL9, ACTB), and myeloid-associated transcripts (C1QA, C1QC, CD163). Overall, stromal transcriptional profiles varied substantially across samples, reflecting differences in histological architecture and RNA capture efficiency.

### 3.3 Cohort-level integration reveals conserved stromal architecture across spatially profiled NET sample

To define shared stromal states across the cohort and minimize sample-specific clustering biases, stromal cell polygons from all eight spatially profiled NET samples were integrated and subjected to unsupervised clustering in the Seurat framework. This resulted in identification of ten transcriptionally distinct clusters (Figure 3A, left). Visualization by sample of origin confirmed that the majority of clusters contained contributions from multiple tumors (Figure 3A, right), demonstrating that cluster identity was largely driven by shared biological programs rather than tumor-specific signatures. However, three clusters (3, 9, and 10), exhibited clear sample predominance and reflected spatially observable tumor-stroma interface regions and therefore cannot be attributed to integration failure. Cluster 3 was largely comprised of cell polygons from NET109. Differential expression analysis combined with Loupe spatial visualization revealed that this cluster localized specifically to the tumor-stroma interface, representing a transcriptionally distinct boundary compartment. Cluster 9 was predominantly composed of NET022 cell polygons and similarly mapped to a tumor-stroma boundary region. Closer inspection of NET022 revealed that tumor cell enriched Space Ranger pipeline generated clusters (Clusters 1, 3, and 4 in Figure 1B and Supplementary Table 4) retained partial neuroendocrine (NE) features, expressing CHGB, CGA, and CALCA, yet simultaneously demonstrated robust expression of adenocarcinoma-associated markers including MUC6, EPCAM, KRT7, and REG1A. Importantly, canonical NE markers CHGA and SYP were absent. This hybrid transcriptional profile indicates that NET022 is indicative of a possible mixed neuroendocrine-non-neuroendocrine neoplasm (MiNEN) phenotype rather than well-differentiated G3 NET. Accordingly, the tumor-stroma interface in NET022 is transcriptionally distinct from the interfaces represented by cluster 3 in other tumors with clear NE differentiation, explaining the sample-specific enrichment of cluster 9. Lastly, cluster 10 was primarily dominated by NET040 and NET111. These were the only PanNET samples containing adjacent exocrine pancreas tissue within the profiled area. Spatial mapping demonstrated that this cluster corresponded to the stroma-exocrine pancreas boundary, accounting for its restricted sample distribution.

**Figure 3.**
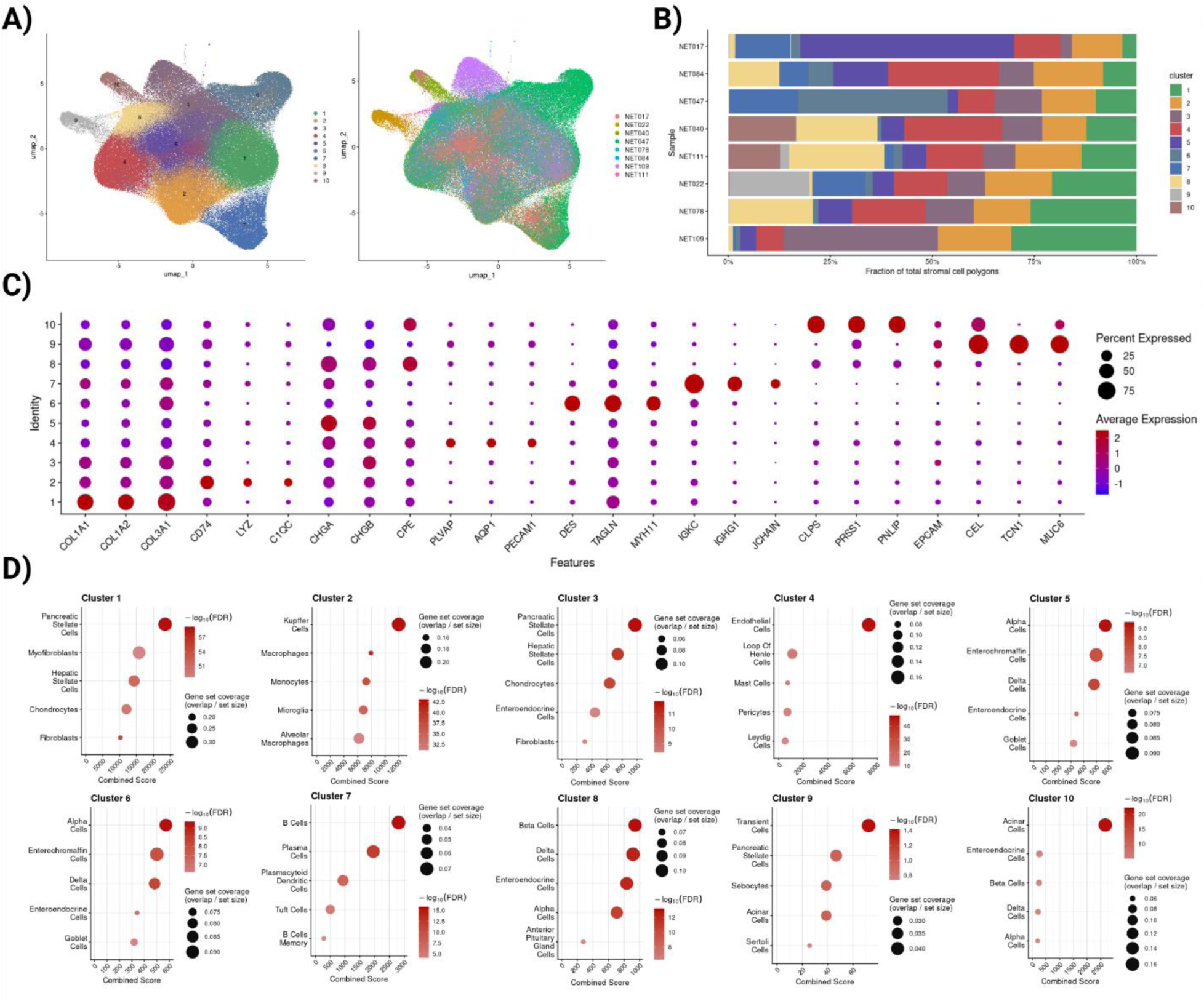
Stromal subset integration. These results demonstrate conserved stromal architecture emerges across eight spatially profiled NET samples. **A)** UMAP representation of integrated stromal cell polygons derived from eight spatial transcriptomics samples. Left: clusters identified by unsupervised integration-based clustering. Right: cells colored by sample of origin, demonstrating effective cross-sample mixing and absence of sample-driven batch segregation. **B)** Relative abundance of integrated stromal clusters across all samples. Stacked bar plots show the fraction of total stromal cell polygons per sample assigned to each cluster, highlighting inter-tumoral heterogeneity in stromal composition while preserving shared stromal states across the cohort. **C)** Canonical marker gene expression across integrated clusters. Dot size represents the percentage of cells expressing a given gene within each cluster, and color intensity reflects average scaled normalized expression. Distinct transcriptional programs support cluster annotation, including extracellular matrix-rich CAF-like populations (COL1A1/COL1A2/COL3A1), myeloid populations (CD74/LYZ/C1QC), endothelial-like cells (PECAM1/AQP1/PLVAP), smooth muscle-like cells (DES/TAGLN/MYH11), B/plasma-like cells (IGKC/IGHG1), residual NET programs (CHGA/CHGB/CPE), and residual glandular epithelial programs (CLPSS, PRSS1, PNLIP, EPCAM, CEL, TCN1, MUC6). **D)** EnrichR cell-type enrichment analysis of cluster-specific marker genes; top 100 DEGs per cluster were selected for enrichment. Dot plots display top 5 enriched cell-types. Dot size indicates gene set coverage (overlap / gene set size), and color intensity represents -log10(FDR).

Despite these boundary-associated exceptions, a conserved stromal framework was evident across the cohort. The relative abundance of tumor/stroma related clusters varied substantially between tumors (Figure 3B) and were concordant with stromal abundance in H&E images. All samples contained extracellular matrix-rich fibroblast-like populations (Figure 3 B-D); however, immune-rich and vascular-associated clusters exhibited marked differences in representation. Desmoplastic tumors (NET109 and NET078) displayed higher fractions of ECM-dominant fibroblast clusters, whereas other tumors showed a higher enrichment of vascular states. These findings indicate that NETs share a common stromal architecture, while individual tumors preferentially expand specific stromal compartments.

Cluster annotation itself was supported by canonical marker gene expression patterns (Figure 3C, Supplementary Table 5). Several clusters were characterized by high expression of fibrillar collagen genes (COL1A1, COL1A2, COL3A1) and additional matrix-associated transcripts, consistent with extracellular matrix-producing CAF-like populations. Distinct non-fibroblast stromal compartments were also resolved, including myeloid populations marked by CD74, LYZ, and complement components (C1QC); endothelial-like cells expressing PECAM1, AQP1, and PLVAP; and smooth muscle-like cells defined by DES, TAGLN, and MYH11. Immune subsets including B/plasma-like cells (IGKC, IGHG1, JCHAIN) were clearly identifiable. Small clusters retained residual tumor-associated neuroendocrine programs (CHGA, CHGB, CPE) or glandular epithelial signatures (CLPS, PRSS1, PNLIP, EPCAM, CEL, TCN1, MUC6), reflecting limited but detectable epithelial carryover despite stringent stromal selection criteria.

To further support cluster identity, we performed cell-type enrichment analysis using the top 100 differentially expressed genes per cluster (Figure 3D). EnrichR-based enrichment confirmed fibroblast/stellate-like signatures in ECM-dominant clusters, macrophage and monocyte signatures in immune clusters, endothelial enrichment in vascular clusters, and smooth muscle/pericyte enrichment in contractile clusters. Boundary-associated clusters demonstrated enrichment for both stromal and residual neuroendocrine or glandular epithelial cell types consistent with their spatial context. The concordance between canonical marker expression and enrichment-based annotation supports robust identification of conserved stromal populations.

### 3.4 CAF subtype identification in shared CAF cluster

To resolve transcriptional heterogeneity within the fibroblast compartment, cells belonging to the shared fibroblast-enriched cluster (Cluster 1) were extracted from the integrated object consisting of stromal compartments from all eight NETs. Rather than re-clustering, subtype identification was performed using literature-derived CAF gene programs and Seurat’s AddModuleScore function (Figure 4A). Four CAF programs were evaluated: myCAF (α-SMA/ECM-related genes), iCAF (IL6/chemokine/NF-κB-associated genes), apCAF (CD74 and MHC class II transcriptional machinery), and csCAF (complement component-enriched genes). Module scores were z-score standardized, and subtype identity was assigned competitively based on relative enrichment within the CAF-core population.

**Figure 4.**
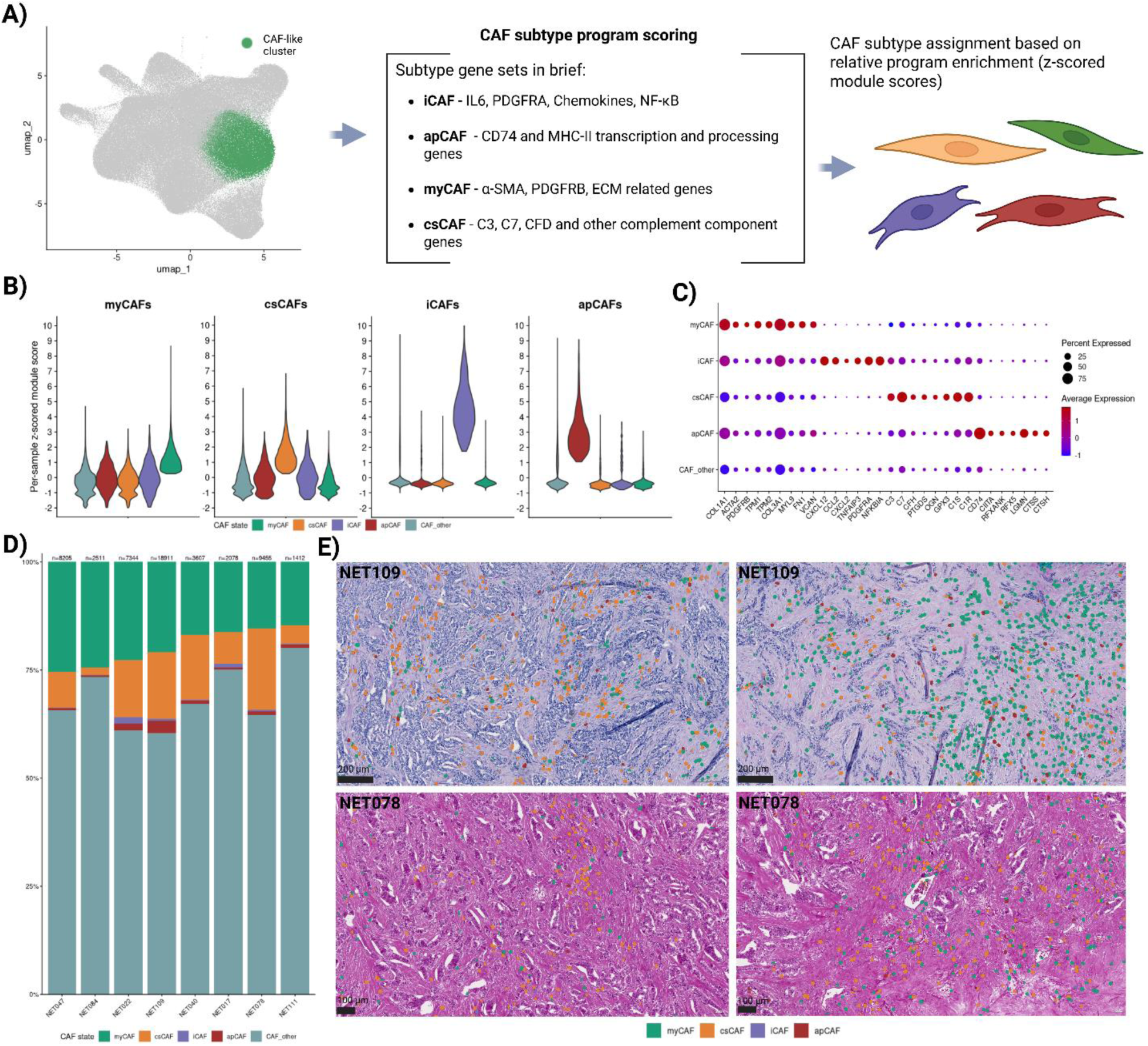
CAF subtypes are conserved across NETs and exhibit distinct spatial organization. (A) Schematic overview of CAF subtype assignment. Cells belonging to CAF-like cluster (CAF-core) were selected for myCAF, iCAF, apCAF, and csCAF program score calculation using literature-derived gene programs and Seurat “AddModuleScore” function. Subtype identity was assigned competitively based on z-score standardized module scores. (B) Violin plots showing per-sample z-score standardized module scores across CAF states. Each subtype exhibits selective enrichment for its corresponding gene program, supporting robustness of subtype classification. (C) Dot plot of representative marker genes for each CAF subtype across all eight samples. Dot size represents the fraction of expressing cells and color indicates average expression level. (D) Relative abundance of CAF subtypes across eight NET samples. Bars represent the fraction of CAF-core cells assigned to each subtype within individual tumors (numbers indicate total CAF-core cells per sample). While myCAFs constitute the dominant population across most tumors, csCAFs represent a substantial complement-enriched fibroblast subset. (E) Spatial distribution of CAF subtypes in representative desmoplastic tumors NET078 and NET109. These samples were selected due to their high stromal content and pronounced CAF-core populations. In tumors lacking a distinct pseudocapsule, csCAFs preferentially localize to tumor-dense stromal interfaces, whereas myCAFs are enriched in collagen-rich desmoplastic compartments, indicating spatially segregated stromal niches.

Violin plots of per-sample standardized module scores demonstrated selective enrichment of each program within its assigned subtype. Cross-enrichment between programs was limited, supporting separation of transcriptional states within the CAF-core population. Subtype assignments were further supported by expression of representative marker genes across all eight tumors (Figure 4C). myCAFs showed elevated expression of COL1A1, COL1A2, TAGLN, and PDGFRB; iCAFs preferentially expressed inflammatory mediators and chemokine-associated transcripts; csCAFs were characterized by complement components including C3, C7, and CFD; and apCAFs demonstrated increased CD74 and additional MHC class II processing related genes. These subtype-specific expression patterns were observed across multiple samples.

Analysis of relative abundance revealed that all four CAF states were present in each tumor, though their proportions varied (Figure 4D). myCAFs constituted the dominant CAF population in most samples. csCAFs represented a substantial complement-enriched subset across several tumors. iCAFs and apCAFs were consistently detected but generally comprised smaller fractions of the CAF-core population. Spatial mapping in representative desmoplastic tumors (NET078 and NET109) demonstrated distinct localization patterns of CAF subtypes (Figure 4E). myCAFs were enriched within collagen-dense stromal regions, whereas csCAFs were preferentially observed at tumor-dense stromal interfaces. These spatial patterns were reproducible across the examined sections.

To further validate CAF state assignments, we performed pseudobulk differential expression analysis (DESeq2) comparing each CAF subtype against all remaining CAF-core cells. myCAFs showed the broadest transcriptional shift, with 41 genes significantly upregulated at FDR < 0.1, of which 30 were not included in the myCAF module-scoring gene set, including additional extracellular matrix and matrix-remodeling transcripts (e.g., SPARC, COL5A1, COL5A2, COL4A2, COL6A1, COL6A2, PLOD2). In contrast, csCAFs displayed a more restricted DEG signature (12 significantly upregulated genes at FDR < 0.1), which largely overlapped with complement program genes used for scoring, with only three additional significant upregulated genes outside the csCAF scoring list (FBLN1, SCARA5, SVEP1). For iCAFs and apCAFs, all significantly upregulated genes overlapped with the respective gene sets used for module scoring, indicating that these subtype-associated DE signals were primarily driven by the scoring program genes in this dataset (Supplementary Table 6). Consistent with these findings, gene set enrichment analysis (GSEA) of Reactome and Hallmark gene sets using genes ranked by the DESeq2 Wald statistic further supported subtype-specific pathway programs (Supplementary Figure 2). myCAFs demonstrated strong enrichment of extracellular matrix organization and collagen-related pathways (including collagen formation/assembly and ECM interactions), as well as Hallmark epithelial-mesenchymal transition, whereas csCAFs showed enrichment for complement cascade/initial triggering of complement and related pathways. iCAF-assigned cells exhibited enrichment for inflammatory/chemokine-associated signaling terms (including chemokine receptor interactions and TNFα/NF-κB- and IFNγ-related gene sets), while apCAF-assigned cells showed enrichment for antigen-presentation-associated signatures (including MHC class II antigen presentation-related terms).

## 4. Discussion

Current classification and diagnostic work-up of neuroendocrine neoplasms are primarily dependent on tumor cell-intrinsic features, including morphology, lineage markers, and proliferative grading as formalized in the 2022 WHO framework (Rindi *et al*, 2022). Increasing evidence, however, indicates that stromal composition and activation state can shape neuroendocrine tumor biology and clinical behavior (Cives *et al*, 2019; Niedra *et al*, 2025; Ye *et al*, 2024). Here, we applied near single-cell resolution spatial transcriptomics across eight treatment-naive primary GEP-NETs to define conserved stromal programs and the *in situ* spatial organization of CAF state programs. Despite modest transcript capture per segmented polygon (median UMI 141; median detected genes 112), Visium HD robustly resolved major tissue compartments and revealed substantial inter-tumoral variation in stromal abundance (8 - 38% of profiled regions; Figure 1B - C). Building on this framework, we inferred the relative abundance and spatial distribution of four CAF programs (myCAF, csCAF, iCAF, and apCAF), providing a cohort-level spatial reference for stromal heterogeneity in NETs beyond single-marker histology.

A central objective of this study was to determine whether a conserved stromal transcriptional framework exists across anatomically distinct GEP-NETs and whether fibroblast activation programs described in carcinoma-type tumors are recapitulated in neuroendocrine neoplasms. Integration of stromal-enriched subsets produced a well-mixed manifold comprising clusters shared across all eight tumors, supporting the presence of conserved stromal transcriptional states rather than sample-driven segregation (Figure 3A). The few sample-predominant clusters localized to spatially defined boundary regions (tumor/stroma interfaces and stroma/exocrine boundaries), indicating niche-restricted states rather than integration artifacts. This cohort-level architecture parallels pan-cancer single-cell studies demonstrating recurrent cross-tumor stromal programs with context-dependent specialization (Li *et al*, 2024; Wang *et al*, 2021; Qian *et al*, 2020). Our spatial integration resolved a distinct mature smooth muscle cell cluster (MYH11, ACTG2, CNN1, SMTN) clearly separated from collagen-rich fibroblast clusters (Figure 3C-D). Given the lower transcript capture inherent to Visium HD, contractile gene expression alone is insufficient to reliably define myCAF identity without careful separation of these mural lineages. Therefore, downstream CAF subtyping was restricted to Cluster 1, which exhibited canonical pan-fibroblast matrix gene expression (COL1A1, COL1A2, COL3A1, MGP, LUM, DCN) consistent with PanglaoDB and Human Protein Atlas annotations (Thul & Lindskog, 2018; Franzén *et al*, 2019).

Pancreatic ductal adenocarcinoma (PDAC) is widely regarded as the reference tumor for CAF biology, as its hallmark desmoplasia has enabled detailed dissection of fibroblast states using both single-cell and spatial transcriptomic approaches (Cui Zhou *et al*, 2022; Chen *et al*, 2026; Yang *et al*, 2023). For this reason, the gene sets used in our study to define myCAF and iCAF programs (Figure 4A; Supplementary Table 1) were primarily curated from PDAC-centered studies and comprehensive CAF reviews (Chen *et al*, 2021b; Tao *et al*, 2025; Melchionna *et al*, 2023; Mori *et al*, 2024; Cui Zhou *et al*, 2022; Chen *et al*, 2021a; Zhang *et al*, 2025; Zhu *et al*, 2025; Elhashani *et al*, 2024; Chen *et al*, 2026; Yang *et al*, 2023). Using these curated gene lists, we observed that the majority of CAFs were assignable to either myCAF or csCAF categories (Figure 4 A - D). The predominance of myCAFs was anticipated, as beyond classical contractile genes (ACTA2, MYL9, TPM1/2), myCAFs are characterized by strong extracellular matrix (ECM) programs and are considered principal effectors of fibrotic matrix deposition within tumor stroma (Hofer *et al*, 2026; Tao *et al*, 2025). This pattern of ECM secretion role was reflected in our dataset at two independent levels. First, pseudobulk differential expression analysis of the myCAF subset demonstrated upregulation of other ECM-associated genes that were outside the list of genes used for myCAF program score calculation (Supplementary Table 6) and subsequent GSEA analysis identified positive enrichment of several ECM related Reactome and Hallmark gene sets (Supplementary Figure 2). Second, in the two most desmoplastic tumors (NET109 and NET078), myCAFs were preferentially localized within collagen-rich stromal regions (Figure 4E), consistent with a structural matrix-maintaining role.

The second most prevalent CAF state - csCAFs, exhibited a distinct spatial pattern and were primarily localized in proximity to tumor cell nests (Figure 4E). A similar spatial association was initially described by Chen et al., who first reported complement-enriched CAFs expressing C3, C7, and related complement components (Chen *et al*, 2021). In our dataset, the csCAF program was robustly supported by strong expressions of C3, C7, C1S, and C1R (Figure 4C), several of which were among the top upregulated genes in the initial fibroblast cluster (Supplementary Table 5). Notably, complement-associated genes can also be expressed by myeloid populations; however, csCAF assignment in our study was restricted to the fibroblast-enriched cluster following stromal integration, minimizing lineage misattribution. Together, these findings suggest that complement-associated fibroblast activation is not restricted to PDAC but may represent a conserved stromal transcriptional program also operative in neuroendocrine tumors.

Compared to myCAFs and csCAFs, evidence supporting inflammatory (iCAF) and antigen-presenting (apCAF) programs was less pronounced. However, this likely reflects technical constraints rather than biological absence. apCAF detection was limited by the Visium HD Human Probe Set v2, which lacks classical MHC class II genes. While CD74 expression alone is insufficient to define apCAFs, its presence within the fibroblast cluster warrants cautious interpretation. Importantly, in our previous GeoMx DSP study (Niedra *et al*, 2025), alpha-SMA-expressing stromal segments exhibited upregulation of CD74 alongside HLA-DRA, HLA-DRB1, and HLA-DPA1 indicative of genuine apCAF presence within NET stroma. Interpretation of iCAFs in NETs may require similar reconsideration. In carcinomas, iCAFs are defined by cytokine/chemokine programs (e.g., IL6, LIF, CXCL family) (Yang *et al*, 2023), yet recent work suggests transcriptional overlap between inflammatory and complement-enriched fibroblast programs, including shared expression of C3, C7, and CFD (Chen *et al*, 2026). Prior scRNA-seq-based PanNET studies (Ye *et al*, 2024; Zhou *et al*, 2021) reported iCAF populations; however, marker sets used to define these clusters included CFD alongside inflammatory genes, which may blur distinctions between chemokine-dominant iCAFs and complement-enriched csCAF-like programs. Given that csCAFs were robustly detected in our cohort, whereas evidence for classical chemokine-dominant iCAF programs was comparatively less prominent, we hypothesize that some previously annotated iCAF-like populations in PanNETs may partially overlap with a dominant complement-enriched csCAF program.

The present findings show striking concordance with our previous Nanostring GeoMx-based study (Niedra et al., 2025), in which α-SMA-enriched stromal segments exhibited gene expression patterns now recognizable as characteristics of three CAF states: myCAFs (ECM-associated genes), apCAFs (MHC class II genes and CD74), and csCAFs (C1R, C1S, COMP, C3), while also lacking in classical chemokine-dominant iCAF signatures. At the time, enrichment of MHC class II and complement-associated genes was conservatively attributed to immune cell contamination, given the compartment-level resolution. However, the present higher-resolution spatial analysis, combined with shared fibroblast cluster-restricted subtype assignment, strongly suggests that these signals represented genuine apCAF and csCAF states that could not be definitively resolved using earlier methodological approaches.

Several limitations should be noted. First, FFPE-based Visium HD segmentation produces lower transcript depth per polygon than fresh scRNA-seq, potentially limiting sensitivity for low-abundance transcripts such as cytokines and MHC-II processing genes relevant to iCAF and apCAF programs. Second, lateral RNA diffusion remains an inherent challenge in spatial transcriptomics, including Visium HD, and may introduce tumor-derived ambient signal into adjacent stromal polygons. While tools such as SpaDiff (Cai *et al*, 2025) and SpotClean (Ni *et al*, 2022) model diffusion in earlier Visium chemistries, they have not been systematically adapted or benchmarked for Visium HD with segmentation-based aggregation. We, to some extent, mitigated this by filtering out cell polygons with high levels of mixed lineage transcriptional patterns and spatial cross-validation, but residual diffusion cannot be fully excluded. Finally, the cohort size (n=8) limits statistical power for subtype prevalence comparisons and precludes correlation with clinical outcomes; metastatic and therapy-exposed lesions were not examined, and NET-specific treatments such as somatostatin analogues (Rogoza *et al*, 2022) may alter stromal transcriptional programs. Despite these constraints, these results extend CAF biology beyond carcinoma paradigms and demonstrate that anatomically distinct GEP-NETs share a conserved stromal transcriptional backbone in which matrix-producing and complement-enriched fibroblast are the dominant states that form spatially segregated niches within NET stroma.

## 5. Data availability

Raw sequencing data (FASTQ files) generated for this study are currently being deposited in the NCBI Sequence Read Archive (SRA) and Gene Expression Omnibus (GEO). Processed GEO data will include spatial gene expression matrices generated using Space Ranger v4.0.1, including segmentation-level filtered feature matrices, spatial metadata, and associated Loupe Browser files. Upon deposition this section will be adjusted with appropriate SRA and GEO reference numbers.

All additional data supporting the findings of this study are available from the corresponding author upon reasonable request.

## 6. Author contributions

HN - data analysis framework development, secondary analyses, data interpretation, manuscript preparation, literature review. MLS - manuscript preparation, literature review. KT - data pre-preparation, cell segmentation. RP and RS - manuscript revisions, statistical analysis verification. JN - patient sample preparation, histological assessment. AO, SV, NS, AG, IK, AP - patient recruitment and clinical data curation. VR - study supervision, manuscript revisions, research team coordination.

## 8. Acknowledgements

This research was also supported within the framework of the European Union’s Recovery and Resilience Mechanism project No. 5.2.1.1.i.0/2/24/I/CFLA/001 “Consolidation of the Latvian Institute of Organic Synthesis and the Latvian Biomedical Research and Study Centre”; specific research project identifier - No. 25/BMC/ZG.

The authors also acknowledge the Latvian Biomedical Research and Study Centre and the Genome Database of the Latvian Population and Pauls Stradins Clinical University Hospital for providing patient’s samples. During the study and manuscript preparation, Marta L. Springe was supported by the framework of the European Union’s Structural Funds Project No.: 1.1.1.8/1/24/I/003; “Strengthening Doctoral Research and Development Capacity at University of Latvia in Smart Specialization Areas”.

